# PP2A^Rts1^ antagonizes Rck2-mediated hyperosmotic stress signaling in yeast

**DOI:** 10.1101/2020.05.30.124925

**Authors:** D. Hollenstein, J. Veis, N. Romanov, G. Gérecová, E. Ogris, M. Hartl, G. Ammerer, W. Reiter

## Abstract

In *Saccharomyces cerevisiae* impairment of protein phosphatase PP2A^Rts1^ leads to temperature and hyperosmotic stress sensitivity, yet the underlying mechanism and the scope of action of the phosphatase in the stress response remain elusive. Using quantitative mass spectrometry-based approaches we have identified a set of putative substrate proteins that show both, hyperosmotic stress- and PP2A^Rts1^-dependent changes in their phosphorylation pattern. A comparative analysis with published MS-shotgun data revealed that the phosphorylation status of many of these sites is regulated by the MAPKAP kinase Rck2, suggesting a node of regulation. Detailed gel mobility shift assays and protein-protein interaction analysis strongly suggest that Rck2 activity is directly regulated by PP2A^Rts1^ via a SLiM B56-family interaction motif, uncovering a previously unknown mechanism of how PP2A influences the response to hyperosmotic stress in Yeast.

## Introduction

The response and adaptation of a cellular system to intra- and extracellular conditions is ensured by signaling networks, commonly mediated by reversible protein phosphorylation. Such networks have been generally conceptualized as being driven by the antagonizing functions of phosphatases and kinases, which tune essential properties of key enzymes by phosphorylation and dephosphorylation. For most signaling events, kinases have been suggested to be the primary activators of downstream targets, conferring signaling specificity by phosphorylating kinase specific protein motifs.

One such a kinase-mediated signaling system is the well-characterized high-osmolarity glycerol (HOG) mitogen-activated kinase (MAPK) pathway in the budding yeast *Saccharomyces cerevisiae* (Hohmann, 2009; Saito and Posas, 2012). Upon elevated extracellular osmolarity, sensory mechanisms of the cell lead to the activation of a signaling cascade, comprising the MAPKK kinases Ssk2, Ssk22, and Ste11 and their substrate MAPK kinase Pbs2. Pbs2. Pbs2 in turn activates the central MAPK Hog1 by dual phosphorylation of Thr^174^ and Tyr^176^. Activated Hog1 orchestrates the effective hyperosmotic stress response by phosphorylating target proteins at specific motifs, characterized by a serine or threonine followed by a proline (S/T-P) (Hohmann, 2009; Saito and Posas, 2012).

We have recently shown that the phosphorylation pattern of more than 200 proteins becomes affected in a Hog1-dependent way upon exposure to hyperosmotic stress. However, the active kinase affects only a limited set of direct substrate sites, phosphorylating S/T-P motifs of roughly 50 target proteins in different cellular compartments (Janschitz et al., 2019; Romanov et al., 2017). The majority of Hog1-affected phosphorylation sites are thus indirectly regulated. One key substrate of Hog1 is the MAPKAPK (MAPK activated protein kinase) Rck2, which mediates a large portion of those secondary phosphorylation events (Bilsland-Marchesan et al., 2000; Teige et al., 2001). In an effort to dissect the signaling mechanism of the HOG pathway, we have therefore proposed Rck2 as a major effector kinase (Romanov et al., 2017).

A widely observed feature of environmental stresses is the transiency of the induced cellular response, consisting of an initiation, a peak and a decline phase. The Hog1 mediated response, for example, peaks at five minutes after mild stress induction and returns to the original state within 20 to 30 minutes (Saito and Posas, 2012). Such a system requires an adequate mechanism to turn off the response effectively; most reports point towards a phosphatase-dependent negative feedback regulation that could ensure that. For instance, phosphatases, such as Ptc1, Ptc2, Ptc3, Ptp2 and Ptp3, have been described to dephosphorylate key residues of Hog1 key residues of Hog1 and thereby inactivate the kinase (Jacoby et al., 1997; Warmka et al., 2001). However, so far it remains unclear whether the inactivation of Rck2 is accomplished by phosphatases as well and if so, which phosphatase could be responsible.

Here we identified protein phosphatase 2A in association with its regulatory subunit Rts1 (PP2A^Rts1^) as a negative regulator of Rck2 activity. The architecture of the multi-protein complex PP2A in *Saccharomyces cerevisiae* is similar to higher eukaryotes. The trimeric PP2A holoenzyme always includes the scaffold subunit A (encoded by *TPD3*) and one of the two catalytic C-subunits Pph21 or Pph22. Additional association with one of the so-called regulatory B-subunits dictates substrate specificity and intracellular localization of the phosphatase. In yeast only two regulatory subunits have been described - Cdc55 and Rts1, belonging to the B and B’ family respectively. Deletion mutants of *RTS1* were characterized as sensitive to temperature and hyperosmotic stress (Evangelista et al., 1996). Yet the underlying mechanisms, particularly in the hyperosmotic stress response, remain elusive.

Applying quantitative shotgun mass spectrometry (MS) we found a substantial overlap between Rts1- and Rck2-dependent phosphorylation events. Detection of *in vivo* interaction between Rts1 and Rck2, as well as its dependence on the SLiM B56-family interaction motif (Hertz et al., 2016; Wang et al., 2016, 2020; Wu et al., 2017) at the C-terminus of Rck2, further strengthens the assumption that Rck2 is indeed a genuine substrate of PP2A^Rts1^. The absence of *RTS1* or mutations of the Rck2 SLiM motif result in a prolonged hyperphosphorylation of Rck2 after stress exposure, arguing for a critical role of PP2A^Rts1^ in the timely de-activation of Rck2 after the termination of Hog1 signaling.

## Results

### PP2A^Rts1^ affects the Rck2 dependent stress phosphorylome

In order to capture the impact of PP2A^Rts1^ on the phospho-proteome of exponentially growing yeast cells, we performed a large-scale SILAC based quantitative MS experiment (Ficarro et al., 2002; Ong et al., 2002) using an *rts1*Δ strain (Figure 1A, Supplementary Table 1). TiO_2_-based phospho-peptide enrichment was combined with strong cation exchange (SCX) fractionation to allow deep phosphoproteome coverage. We performed five independent biological replicate experiments and analyzed a total of 35 LC-MS (liquid chromatography-mass spectrometry) runs. We observed that ~95% of all protein SILAC ratios were clustered between 0.5 and 2, hence no protein normalization was performed (Supplementary Figure 1A). SILAC ratios from peptides containing the same set of phosphorylated residues were grouped and defined as phosphorylation site groups (further referred to as phosphorylation sites). Measures, such as the reproducibility of the SILAC ratios for the phosphorylation sites (Supplementary Figure 1B, average *R* ~ 0.73), demonstrate the quality and consistency of the MS datasets, and that they can be reliably compared with other published data. Overall our analysis resulted in 8.611 quantified phosphorylation sites. Phosphorylation sites displaying a more than two-fold change were considered to be affected by the deletion of *RTS1*. Most of the affected sites tend to become increased (fold change ≥ 2) in phosphorylation rather than decreased (fold change ≤ 0.5), which is the expected effect of phosphatase inactivation. Specifically, 84.9% of the sites were unchanged, 11.3% had an increase and 3.9% a decrease in phosphorylation frequency (Figure 1B). To confirm that our experimental setup adequately reflected the anticipated effects of an *RTS1* deletion, we compared the MS results with previously reported hallmarks of PP2A^Rts1^ dependent signaling. Indeed, several described substrates of PP2A^Rts1^ showed increased phosphorylation, such as Ace2 (Parnell et al., 2014), Swi6 (Artiles et al., 2009), Kin4 (Chan and Amon, 2009; Zuzuarregui et al., 2012) and the inhibitory Tyr^19^ of Cdk1/Cdc28 (Harvey et al., 2011) (Supplementary Table 2).

**Figure 1.**
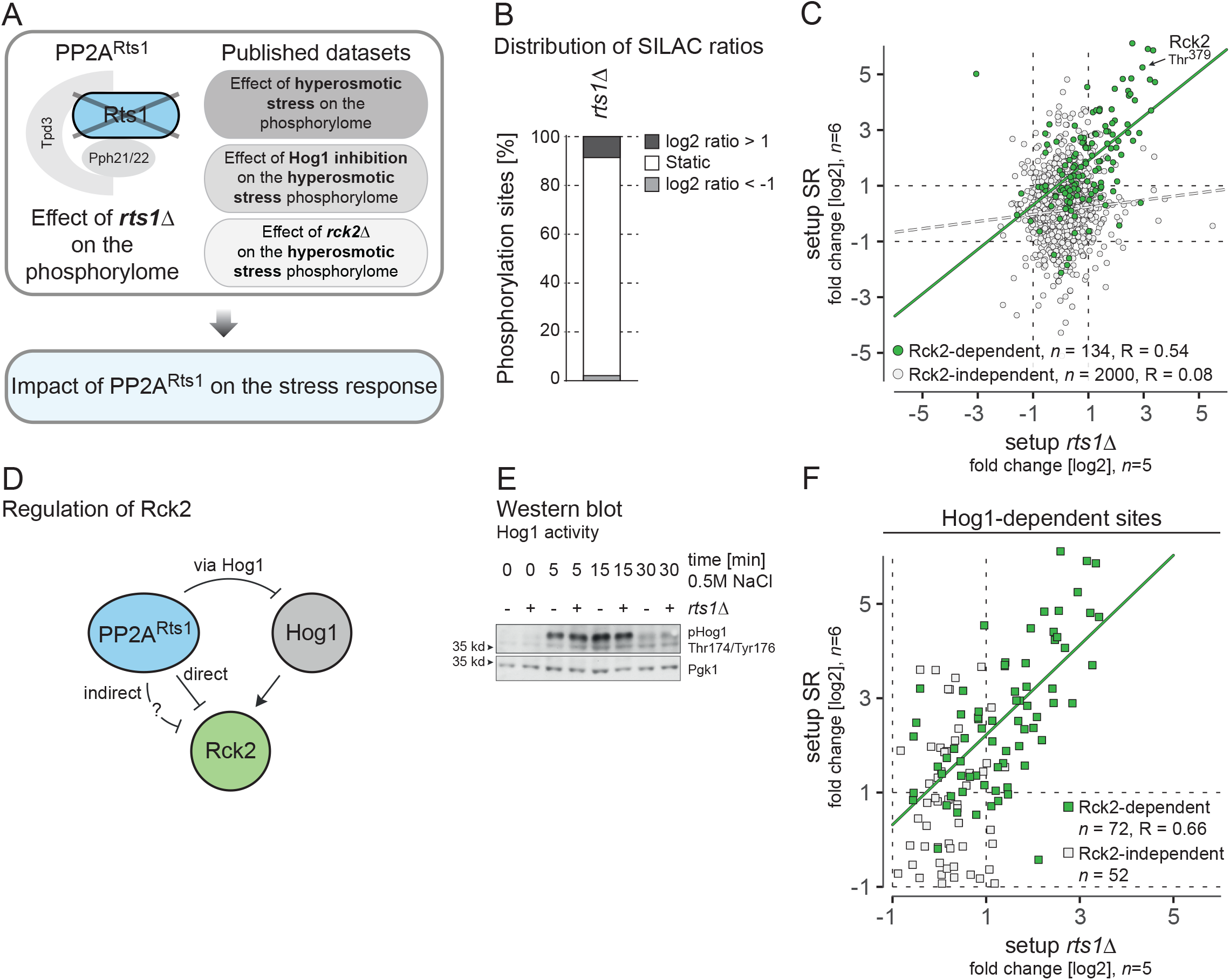
Deletion of Rts1 results in hyperactivation of the MAPKAPK Rck2 but not of the MAPK Hog1. (A) Quantitative MS-based phospho-proteomic experiments used in this study. Experiments performed in this study encompass *RTS1* deletion strains (*n* = 5); the derived data was compared with the published MS dataset that captured the early response to hyperosmotic stress (Romanov et al., 2017). All SILAC ratios represent knockout versus wild type or stressed versus unstressed. (B) Increased phosphorylation in contrast to static sites and de-phosphorylation observed in *rts1*Δ MS-shotgun experiments. (C) Scatter plot displaying log2 SILAC ratios of individual phosphorylation sites (grey) in the experiments SR on the *x*-axis and setup *rts1*Δ on the *y*-axis. Dots in green represent phosphorylation sites that are known to be targets of Rck2 (Setup SR *rck2*Δ fold change ≤ 0.5) (Romanov et al., 2017). (D) Schematic illustration of potential regulatory mechanism of Rck2 by PP2A^Rts1^. (E) Western blot showing Hog1 kinase activation loop phosphorylation in exponentially growing cells (0 minutes treatment) and cells treated for five minutes with 0.5 M NaCl. Deletion of *RTS1* has no effect on Hog1 double-phosphorylation of Thr^174^ and Tyr^176^. pHog1: Phospho-p38 antibody, Pgk1: Pgk1 antibody - loading control. (F) Scatter plot displaying log2 SILAC ratios individual Hog1-dependent phosphorylation sites (Setup SR Hog1as fold change ≤ 0.5) in the experiments “setup *rts1*Δ*”* on the *x*-axis and “SR” on the *y*-axis. Phosphorylation sites are classified according to the observed behavior in the “SR *rck2*Δ” experiment (Romanov et al., 2017). Rck2-dependent phosphorylation sites (Setup SR *rck2*Δ fold change ≤ 0.5) are shown in green, and Rck2-independent phosphorylation sites (Setup SR *rck2*Δ fold change > 0.5) as grey squares. SR: stress response

To identify phosphorylation sites that mediate the influence of PP2A^Rts1^ on the hyperosmotic stress response we integrated the *rts1*Δ MS results with previously published large-scale quantitative MS-based proteomic datasets, which characterized the phosphoproteomic state in response to hyperosmotic stress (further referred to as setup SR [stress response]) (Romanov et al., 2017) (Figure 1A). We could not detect any strong correlation between fold changes when comparing stress- and *rts1*Δ-affected phosphorylation sites, suggesting that the Rts1 impact on the hyperosmotic stress response might be restricted to a small sub-regulon. Interestingly, we found Rck2 Thr^379^ - the key regulatory residue in the activation loop of the MAPKAP kinase - to be affected by the absence of PP2A^Rts1^ (fold change ~ 7.76 in *rts1*Δ). Key activating residues of four other kinases that were recovered in the *rts1*Δ experiment were not affected by *RTS1* deletion. Given that we previously described the Rck2 sub-regulon of the HOG response, we focused on Rck2-dependent phosphorylation sites and analyzed their phosphorylation state in the *rts1*Δ dataset. Remarkably, fold changes of Rck2-dependent phosphorylation sites were significantly correlated between the stress response and the *rts1*Δ datasets, suggesting that PP2A^Rts1^ exerts its impact on the hyperosmotic stress response specifically through Rck2 (Figure 1C). Moreover, a substantial part of the Rck2- and hyperosmotic stress-affected set of phosphorylation sites displayed strongly increased phosphorylation in *rts1*Δ cells (57%, fold change ≥ 2). We therefore assume that even in the absence of hyperosmotic stress, Rck2 becomes activated in PP2A^Rts1^-deficient cells. This could be due to the absence of direct dephosphorylation of Rck2 by PPA2^Rts1^ or indirectly, via elevated activity of Hog1, or due to another, yet unknown regulatory mechanism (Figure 1D).

Since Thr^379^ of Rck2 is a well-established substrate of the MAPK Hog1 (Teige et al., 2001), we investigated to which extent Hog1 activity is affected by PP2A^Rts1^ inactivation. Thr^174^ and Tyr^176^, the key residues of the Hog1 activation loop, did not show any increase in phosphorylation in *rts1*Δ cells. This was both evident from the MS-data (Figure 1C), as well as separate Hog1 activity measurements using an antibody specifically recognizing Thr^174^/Tyr^176^ double phosphorylation of Hog1 (Figure 1E). Moreover, Hog1-dependent phosphorylation sites that are not regulated by Rck2 were also not affected by *RTS1* deletion. On the other hand, phosphorylation sites of the Rck2 sub-regulon were indeed strongly induced in the *rts1*Δ dataset (Figure 1F). Thus, we concluded that PP2A^Rts1^ regulates Rck2 activity not via Hog1, but either directly or via some other unknown kinase or phosphatase.

### PP2A^Rts1^ interacts with Rck2 via a SLiM motif

To test whether PP2A^Rts1^ directly interacts with Rck2 *in vivo* we performed an M-track protein-proximity experiment (Brezovich et al., 2015; Zuzuarregui et al., 2012). As prey, Rck2 was tagged with histone H3-hemagglutinin (HA) tag fused to protein A (protA-H3-HA) and co-expressed with Rts1 that was fused to the enzymatic domain of the murine histone lysine methyltransferase SUV39 (HKMT-myc) serving as bait. Upon close proximity of bait- and prey-protein the protA-H3-HA tag becomes irreversibly tri-methylated on lysine 9 of the histone H3 (me3K9H3). We observed strong proximity signals between Rts1-HKMT-myc and Rck2-protA-H3-HA, but no methylation when Rts1 was not tagged, confirming that Rts1 and Rck2 interact *in vivo* (Figure 2A).

**Figure 2.**
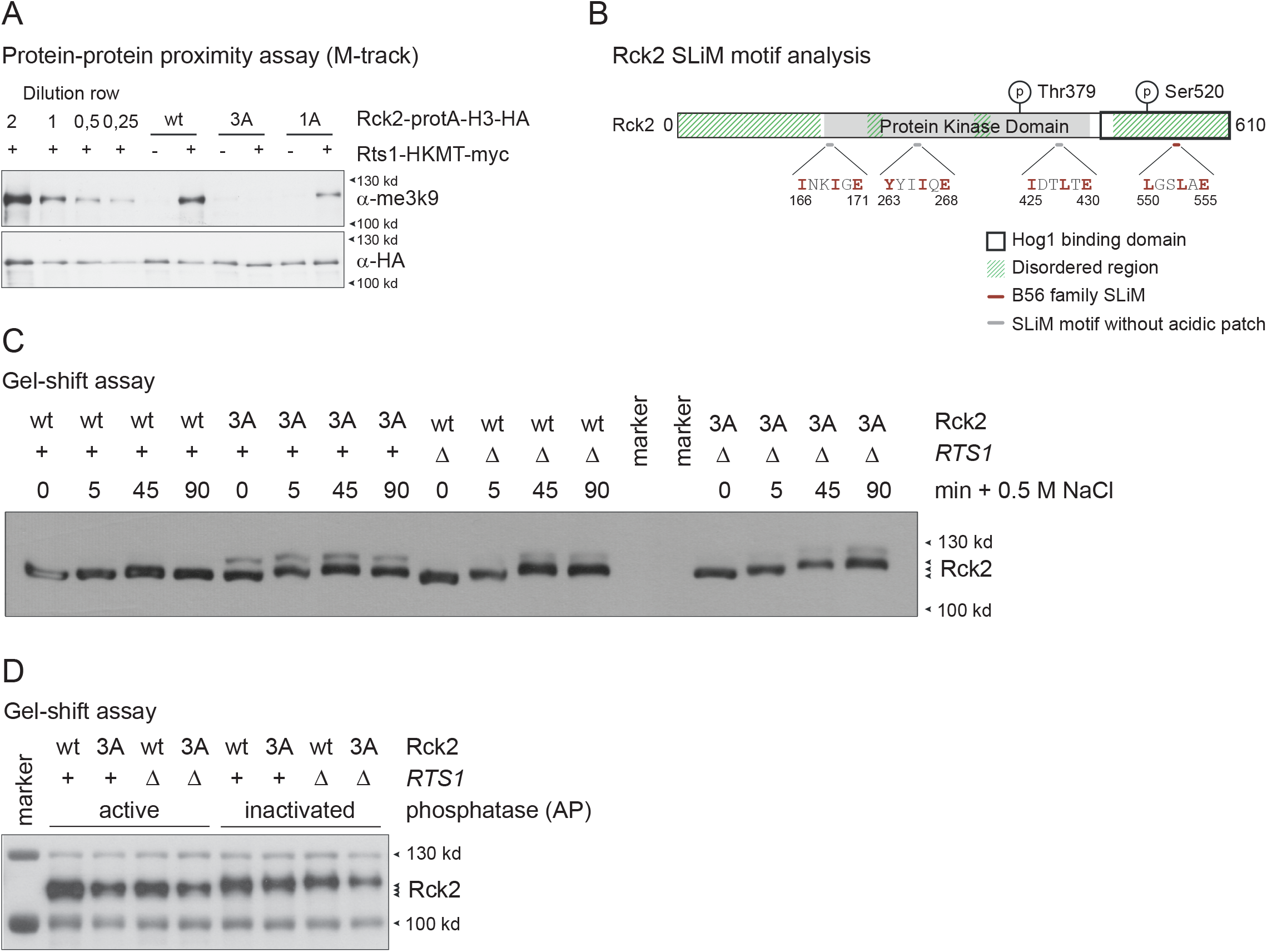
PP2A^Rts1^ interacts with Rck2 via a SLiM motif. (A) Western blot showing M-track results obtained for Rts1-HKMT-myc and Rck2-protA-H3-HA interaction. Background signal intensity was defined using a strain expressing Rck2-protA-H3-HA and an untagged wildtype allele of Rts1. M-track proximity signals were obtained from exponentially growing cells. Detection of M-track signal with H3K9 trimethylation antibody. Control: HA antibody. Much less or no signal was detected between Rts1-HKMT-myc and mutant versions of Rck2-protA-H3-HA, namely 1A and 3A. (B) Schematic showing positions of predicted SLiM motifs in the protein sequence of Rck2. Only the motif at position 550-561 is followed by a sequence of acidic residues, and lies in a predicted disordered region of the protein. Amino acid positions of motifs are indicated. (C) Phosphorylation of Rck2 after hyperosmotic stress depends on PP2A^Rts1^ and the Rck2-SLiM motif: Western blot showing phosphorylation-shifts of Rck2 purified from different strains that were exposed to 0, 5, 45, and 90 minutes hyperosmotic stress, respectively. (D) The 45 minutes stress samples from (C) were treated with active or heat inactivated alkaline phosphatase before analysis by western blotting. To facilitate protein alignment, samples were spiked with a prestained protein ladder, and the 100 and 130 kDa proteins of the protein ladder were visualized using the antiblue-antibody. AP: alkaline phosphatase; wt: wild type

It has been previously demonstrated that substrate-binding of B56, the human ortholog of Rts1, requires the presence of a fully functional small linear binding motif (SLiM) (Hertz et al., 2016; Suijkerbuijk et al., 2012). An example for such an interaction has also been described for yeast Rts1 that was able to interact with the mammalian homolog of the spindle checkpoint protein Mad3 in mammalian cells (Hertz et al., 2016). Therefore, we reasoned that the interaction between PP2A^Rts1^ and Rck2 might also be mediated via a SLiM binding motif. We identified four potential LxxIxE motifs in the protein sequence of Rck2, namely INKIGE (amino acid 166 to 171), YYIIQE (263-268), IDTLTE (425-430) and LGSLAE (550-555) (Figure 2B). Functional SLiM motifs are found within intrinsically disordered regions (Ren et al., 2008). We therefore utilized the SPOT-Disorder algorithm (Hanson et al., 2017) to predict disordered regions in the Rck2 protein sequence. Of the three potential LxxIxE motifs, only LGSLAE (550-555) was located within a putative disordered region. This motif contains a Leucine at position 1 and 3, which was predicted to confer high binding affinity to Rts1. In addition, the motif is followed by an acidic patch that contains five acidic residues. Taken together, our computational analyses suggested that residues 550 to 555 of Rck2 constitute an optimal LxxIxE motif that enables recruitment of PP2A^Rts1^. It should be noted that other previously described targets of PP2A^Rts1^, such as Ace2, Cdc28, Kin4 and Swi6, also contain putative SLiM motifs (Supplementary Figure 2A). To test whether the interaction between PP2A^Rts1^ and Rck2 requires this functional SLiM, we created point mutations within the C-terminus of Rck2, namely Rck2-1A (E555A) and Rck2-3A (L550A/L553A/E555A). We tested these mutant alleles for the ability to bind PP2A^Rts1^ *in vivo* using M-track assays (Figure 2A, see lanes labeled with *3A* and *1A*). Contrary to what we observed for the Rck2 wildtype, the interaction signal fades significantly for 1A, and completely disappears for the 3A-mutant. Our results thus demonstrate that PP2A^Rts1^ directly interacts with Rck2 via a SLiM motif in the C-terminus of Rck2.

### Dephosphorylation of Rck2 after hyperosmotic stress depends on PP2A^Rts1^ and the SLiM motif

After establishing that Rck2 indeed directly interacts with PP2A^Rts1^, we sought to understand whether Rck2 represents an actual substrate of the phosphatase. For this purpose, we first examined whether the phosphorylation status of Rck2 is affected in cells expressing mutant alleles of the SLiM motif. Using gel mobility shift assays we could observe that the migration of the Flag-tagged wildtype and mutant Rck2 before and after exposure to hyperosmotic stress (0.5 M NaCl for 5, 45 and 90 minutes) differed substantially (Figure 2C and Supplementary Figure 2B). Treatment with alkaline phosphatase abolished the shifts, confirming that the observed mobility shifts are indeed caused by phosphorylation (Figure 2D and Supplementary Figure 2C). The phosphorylation pattern of Rck2 varied according to the presence of PP2A^Rts1^, but also according to whether Rck2 carried the mutations in the SLiM motif. More specifically, we observed an upshift on the gel for the mutated version of Rck2. Notably, phosphorylation-induced shifts in gel-mobility have been mostly associated with the key regulatory site of Rck2 (Teige et al., 2001). We thus conclude that the dephosphorylation of Rck2 after termination of Hog1 signaling is indeed mediated by direct interaction of PP2A^Rts1^ with Rck2 via its SLiM motif (Figure 2C and 2D).

## Discussion

Several aspects of the functional role of PP2A^Rts1^ in the environmental stress response of *Saccharomyces cerevisiae* remain elusive. It is neither known whether the trimeric phosphatase complex exerts (de-)phosphorylation in a direct or indirect manner, nor how wide-ranging its impact actually is. The investigation of phosphatases is substantially impeded by the transient nature of interactions between phosphatases and their substrates, complicating a clear-cut inference on underlying mechanisms. In the presented work, we make use of a number of experimental methods and techniques to understand the role of PP2A^Rts1^ in the hyperosmotic stress response.

By means of SILAC-based quantitative MS-analysis of the phosphorylome in *rts1*Δ cells, we observed increased phosphorylation at the activation loop of the MAPKAP kinase Rck2, which we recently identified as a central effector kinase downstream of Hog1, controlling the majority of secondary Hog1 phosphorylation events (Romanov et al., 2017). While inhibition of Hog1 activity during termination of the stress response involves negative feedback loops, which include multiple tyrosine and serine/threonine phosphatases (English et al., 2015; Maeda et al., 1993; Wurgler-Murphy et al., 1997), it is not known how Rck2 signaling is downregulated. Our speculation on a potential connection between PP2A^Rts1^ and Rck2 was further substantiated by the presence of a SLiM motif in a disordered region at the C-terminus of the Rck2-kinase. Such a motif would provide the means for a direct interaction between the phosphatase and its potential substrate, as it has been demonstrated for B56, the human paralog of Rts1 (Hertz et al., 2016). We thus followed up on our hypothesis that PP2A^Rts1^ might directly regulate Rck2 activity in response to hyperosmotic stress.

Our experiments demonstrate that increased Rck2 activity resulting from an *RTS1* deletion is not caused by misregulation of Hog1 activity. Instead, the positive result of the protein proximity assay along with the presence of an LxxIxE SLiM motif in the Rck2 protein sequence strongly favors a model in which PP2A^Rts1^ negatively regulates Rck2 activity by dephosphorylation of the key regulatory residue Thr^379^. Our results further suggest PP2A^Rts1^ to be a critical component required for the adequate shut-down of Rck2 activity once the hyperosmotic stress response is terminated. At the same time the *rts1*Δ MS results indicate that the phosphatase keeps the basal phosphorylation status of the kinase at a low level. Given these insights, we speculate that the primary role of PP2A^Rts1^ lies most likely in the termination of stress signaling pathways, which could also be relevant for the function of PP2A^Rts1^ in other cellular processes (Zapata et al., 2014).

Our findings remain confined to the specific interplay of PP2A^Rts1^ and Rck2, and do not clarify whether proteins downstream of Rck2 signaling are direct PP2A^Rts1^ substrates as well. Adequate changes in their phosphorylation status and activity would be eminently needed to ensure proper termination of the hyperosmotic stress response.

We identified putative SLiM motifs in the protein sequences of four known targets of PP2A^Rts1^. It is interesting to speculate that SLiM-mediated interaction might represent a widespread mechanism for phosphatase interaction in signaling pathways in general (Brautigan and Shenolikar, 2018; Tompa et al., 2014). Given that deletion of *RTS1* results in activation of Rck2 and, thereby, indirectly in increased phosphorylation of the whole Rck2 regulon, putative substrates of the phosphatase should be evaluated critically. The presence of a functional SLiM motif and the dependence of phosphorylation changes on this motif might help to identify direct substrates of the phosphatase.

Another key question that remains to be answered concerns the actual regulation of PP2A^Rts1^ activity. Rts1 is a highly phosphorylated protein (Alcaide-Gavilán et al., 2018; Shu et al., 1997), however, the consequences of phosphorylation on its activity remain uncharacterized. Reports from mammalian cells indicate that some phosphorylation might be activating (Xu and Williams, 2000), whereas others might have inhibitory effects (Letourneux et al., 2006). Future studies are necessary to determine whether PP2A^Rts1^ is constitutively active or somehow regulated by stress signaling. The hyperosmotic stress response might in this regard serve as an ideal starting point for dissecting regulatory mechanisms that control the activity of PP2A^Rts1^.

## Supplementary Figure Legends

**Supplementary Figure 1.**
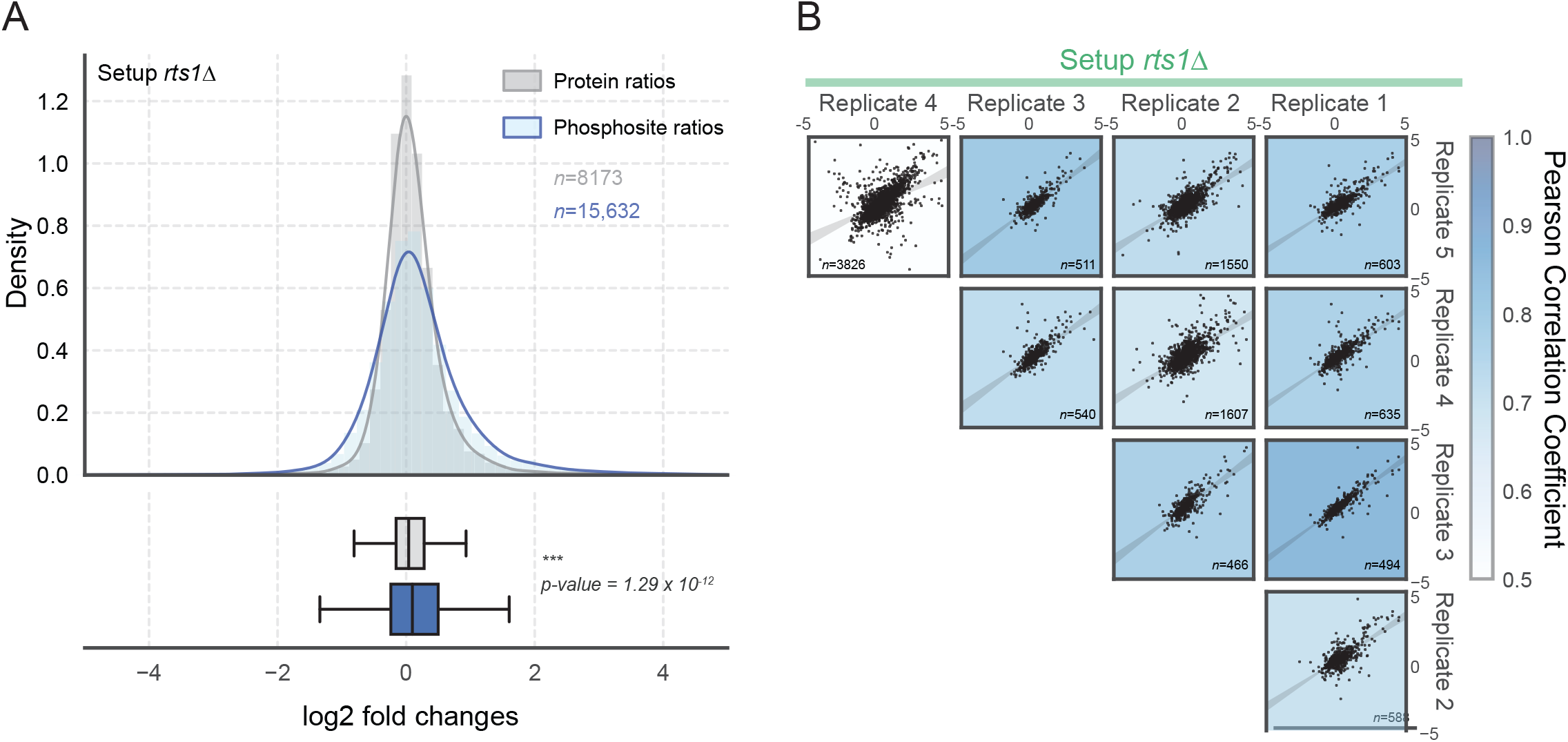
(A) Protein abundance is not affected by deletion of *RTS1*. Histograms and boxplots illustrate SILAC ratios of unphosphorylated (grey) and phosphorylated peptides (blue) from setup *rts1*Δ. *P*-values were calculated using a *t*-test. (B) SILAC ratios are reproducible across biological replicates (illustrated for setup *rts1*Δ). For each scatter plot the background color reflects the underlying Pearson correlation coefficient. The number of data points is shown in each scatter (*n*), and the shaded grey bands indicate the regression estimate at the confidence interval of 0.9.

**Supplementary Figure 2.**
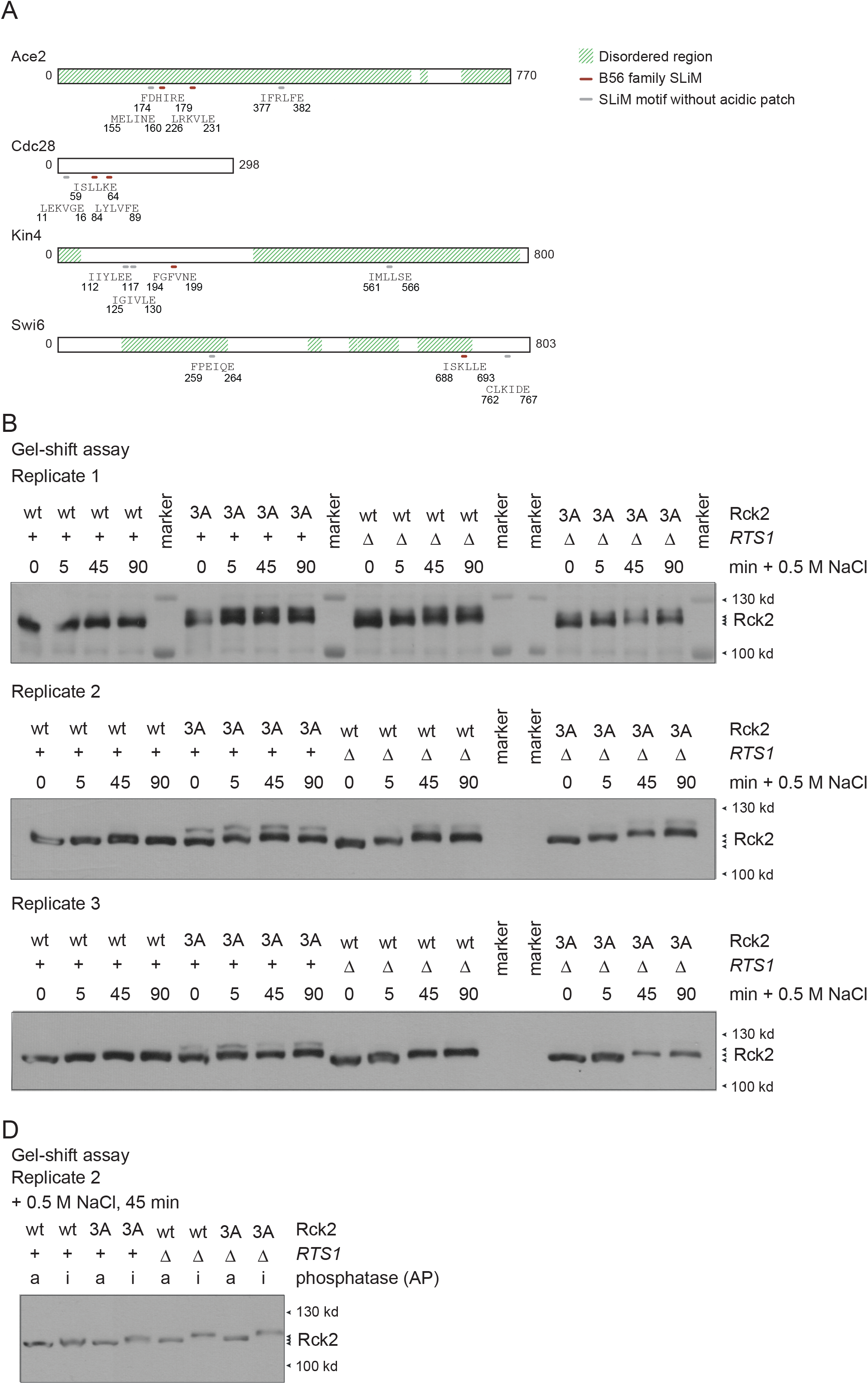
(A) Schematic showing positions of predicted SLiM motifs and disordered regions of Ace2, Cdc28, Kin4 and Swi6. Amino acid positions of motifs are indicated. (B) Replicate data for Figure 2C: Western blot showing phosphorylation-shifts of Rck2 purified from different strains that were exposed to 0, 5, 45, and 90 minutes hyperosmotic stress, respectively. (C) Second replicate for Figure 2D: The 45 minutes stress samples from (B) were treated with active or heat inactivated alkaline phosphatase (AP) before analysis by western blotting. a: treated with active alkaline phosphatase, i: treated with heat-inactivated alkaline phosphatase.

## Supplementary Table Legends

**Supplementary Table 1.** Overview on the experimental setup of the MS experiments containing details on experiment name, experimental condition, control condition, strains used, and SILAC labeling.

**Supplementary Table S2.** Summary of quantified phosphorylation sites over MS shotgun experiments of our study, and previous studies (four experimental conditions in total). The table includes SILAC ratios of quantified phosphorylation sites in different conditions.

**Supplementary Table S3.** Strains used in this study.

**Supplementary Table S4.** Detailed description of mass spectrometer acquisition settings of individual raw files.

## Materials & Methods

### Yeast strains and plasmids

*Saccharomyces cerevisiae* strains used in this study are listed in Supplementary Table 3. Standard methods were applied to perform genetic manipulations of plasmids and yeast strains. Yeast strains used for MS experiments and M-track assays were derived from WR209, a W303 SILAC wild type strain (Romanov et al., 2017). All other yeast strains are derivatives of W303-1A.

### Growth conditions, SILAC labeling and harvesting

Yeast cells were grown shaking (200 rpm) at 30 °C for at least ten generations until mid-log phase (OD_600 nm_ ~1). For M-Track experiments and gene expression analysis yeast cultures were grown in 50 ml rich medium (YPD; 1% yeast extract, 2% peptone, 2% glucose), harvested by centrifugation (2 × 1.5 min, 3G) and frozen in liquid N_2_. For MS experiments SILAC labeling (Gruhler et al., 2005; Ong et al., 2002) was performed as follows: Yeast cultures were grown in 50 ml synthetic medium (0.17% yeast nitrogen base, 0.5% ammonium sulfate, 2% glucose, amino acids as required) supplemented either with heavy or light labeled arginine and lysine (0.05 mg/ml of L-arginine-HCl and 0.05 mg/ml of L-lysine-2HCl, Euriso-top), harvested by filtration (Protran 0.45 μm nitrocellulose membrane, Amersham) and frozen in liquid N_2_. For each experiment, heavy and light labeled cultures were grown in parallel and frozen pellets were united after harvesting. Details on the setup of MS experiments are provided in Supplementary Table 1.

### Protein extraction and enzymatic digestion for MS experiments

Proteins were extracted using TRIzol (Invitrogen) (Reiter et al., 2012), or (for experiments 08_rts1_1 and 08_rts1_2) using a TCA based protocol. TCA extraction was performed as follows: cells were resuspended in 8 M urea, 50 mM Tris pH 8.0 and disrupted by beat beating using a Fast Prep (3 cycles: 45 s, power level 5.5). Insoluble material was removed by centrifugation. Proteins were extracted by addition of ice-cold TCA (15% final concentration), followed by an incubation for 60 minutes on ice. Proteins were centrifuged (12,000 x g, 5min, 4 °C), washed in 15% TCA and acetone and shortly dried. Protein pellets were resuspended in 50 mM ammonium bicarbonate (ABC) buffer containing 8 M urea. Protein concentration (2-3 mg/ml) was determined by Bradford protein assay (Bio Rad), using bovine serum albumin to create a standard curve. Protein samples were diluted to 50 mM ABC, 6 M Urea by using 50 mM ABC. Disulfide bridges were reduced by adding dithiothreitol (DTT), using a DTT to protein ratio of 1:50, and samples were incubated for 30 minutes at 56 °C. Cysteines were alkylated by adding iodoacetamide (IAA), using an IAA to protein ratio of 1:10, and samples were incubated for 30 minutes in the dark at room temperature. Remaining IAA was quenched by adding DTT, using a DTT to protein ratio of 1:100. Proteins were digested with LysC (Roche) for 2 hours at 30 °C, using a LysC to protein ratio of 1:100. Protein samples were diluted to 50 mM ABC 0.6 M Urea by using 50 mM ABC. Proteins were digested with trypsin (Trypsin Gold, Promega) overnight at 37 °C, using a trypsin to protein ratio of 1:60. The overnight digest was stopped by adding 100% trifluoroacetic acid (TFA) to a final concentration of 1%. Resulting peptide samples were desalted using Strata-X reversed phase polymeric solid phase extraction cartridges Phenomenex, 200 mg), and eluted by addition of 70% acetonitrile (ACN) 0.1% formic acid (FA). An aliquot of ~ 1μg protein extract was taken, diluted with 0.1% TFA to an ACN concentration below 2% and subjected to MS analysis. Peptide samples were snap-frozen in liquid nitrogen, lyophilized and stored at −80 °C.

### Phosphopeptide isolation and fractionation

Phosphopeptides were enriched using TiO_2_ (Titansphere bulk media, 5 micron, GL Science). The amount of TiO_2_ resin was adjusted to the peptide concentration (1.25 mg of TiO_2_ / 3.5 mg yeast protein extract). TiO_2_ resin was washed with 50% Methanol, H_2_O and equilibrated with TiO_2_ loading buffer (0.8 M phtalic acid, 80% ACN, 0.1% TFA). Dried peptide samples were dissolved in 100 μl TiO_2_ loading buffer and incubated for 1 hour with 1mg TiO_2_ resin per 2.8 mg protein extract. The TiO_2_ resin were transferred to a Mobicol spin column and washed with 2 × 250 μl TiO_2_ loading buffer, 2 × 250 μl 80% ACN 0.1% TFA, 2 × 250 μl 1% ACN 0.1% TFA. Bound phosphopeptides were eluted by addition of 2 × 150 μl 0.3 M ammonium hydroxide and acidified to pH 2.5 by addition of 10% TFA. Phosphopeptide samples were desalted using C18 Sep-Pak cartridges (Waters), vacuum dried and stored at −80 °C. Phosphopeptides were fractionated offline by strong cation exchange chromatography (SCX), using 1 ml Resource S column (GE healthcare) connected to a nano-HPLC (Ultimate 3000, Thermo Fisher Scientific). Briefly, samples were injected using SCX Buffer A (5 mM NaH_2_PO_4_, 30% acetonitrile (ACN), pH 2.7). Peptides bound to the column were separated by a linear gradient of sodium chloride in SCX buffer A. Based upon UV measurements, fractions containing low amounts of peptide were pooled which resulted in a total of 12 fractions (fractions were collected every minute and then pooled). Each elution sample was adjusted by TFA to pH 2-3 for subsequent desalting (Rappsilber et al., 2007) and mass spectrometry measurement.

### Mass spectrometry measurements

LC-MS/MS analysis was performed on an UltiMate 3000 Dual LC nano-HPLC System (Dionex, Thermo Fisher Scientific), containing both a trapping column for peptide concentration (PepMap C18, 5 × 0.3 mm, 5 μm particle size) and an analytical column (PepMap C18, 500 × 0.075 μm, 2 μm particle size, Thermo Fisher Scientific), coupled to a Linear Trap Quadrupole Orbitrap Velos (with CID, collision-induced dissociation mode; or ETD, electron-transfer dissociation) mass spectrometer (Thermo Fisher) or a Q Exactive HF Orbitrap (with HCD, higher-energy collisional dissociation mode) mass spectrometer (Thermo Fisher) via a Proxeon nanospray flex ion source (Thermo Fisher).

For peptide separation on the HPLC the concentration of organic solvent (acetonitrile) was increased from 2.5% to 40% in 0.1% formic acid at a flow rate of 275 nl/min, using different gradient times. For acquisition of MS2 spectra the instruments were operated in a data-dependent mode with dynamic exclusion enabled. A detailed description of the acquisition settings for individual raw files is listed in Supplementary Table 4.

### Mass spectrometry data analysis with MaxQuant

Raw MS data was analyzed using MaxQuant (Cox and Mann, 2008) software version 1.5.2.8 (global proteome experiments), using default parameters with the following modifications. MS2 spectra were searched against a protein database from the SGD (Saccharomyces Genome Database, www.yeastgenome.org, version 3^rd^ February, 2011) containing 6,717 entries, concatenated with a database of 248 common laboratory contaminants (provided with MaxQuant). Hence, the option to include contaminants was deactivated. Enzyme specificity was set to “Trypsin/P” (allowing cleavage after proline), the minimal peptide length was set to 6 and the maximum number of missed cleavages was set to 2. The option “I = L” was activated to treat the amino acids leucine and isoleucine as indistinguishable. The minimum peptide length was set to 6. Carbamidomethylation of cysteine was defined as fixed modification. “Acetyl (Protein N-term)”, “Deamidation (NQ)”, “Oxidation (M)” and “Phospho (STY)” were set as variable modifications. A maximum of 6 variable modifications per peptide was allowed. For MS measurements of samples prior to phosphopeptide enrichment “Phospho (STY)” was not used as a variable modification. For SILAC quantification “multiplicity” was set to 2, “Arg6” and “Lys6” were specified as heavy labels, “Requantify” and “Match between runs” were enabled.

### Calculation of phosphorylation site SILAC ratios

For calculation and normalization of phosphorylation site SILAC ratios in-house Python scripts were used. All data was extracted from MaxQuant evidence tables. SILAC ratios (heavy to light) were extracted from the column “Ratio H/L”, log2 transformed and, if necessary, inverted (see Supplementary Table 1). SILAC ratios were corrected for differences in the amount of heavy-labeled and light-labeled cells. In addition, proline containing peptides were corrected for signal loss caused by the conversion of heavy-labeled arginine to heavy-labeled proline (Ong et al., 2003). Normalization factors were calculated independently for each replicate and experiment. For calculation of normalization factors only unphosphorylated peptides were considered. First, the average log2 ratio of peptides, not containing proline, was calculated and subtracted from the log2 ratios of individual phosphorylated and unphosphorylated peptides. Second, a proline-conversion factor was calculated as the average log2 ratio of unphosphorylated peptides containing exactly one proline and the log2 ratio (divided by two) of peptides containing two prolines. For each phosphorylated and unphosphorylated peptide the proline-conversion factor was multiplied by the number of prolines present in the peptide sequence and subtracted from the log2 ratio. An isoform phosphorylation site probability was calculated by multiplying the highest individual phosphorylation site probabilities. Peptides with an isoform probability below 70% were discarded. To facilitate interpretation of protein phosphorylation sites, phosphopeptides were grouped into “phosphorylation sites” containing the same set of phosphorylated protein residues, regardless of potential missed cleavages or additional modifications such as oxidation. The SILAC log2 ratio of individual “phosphorylation sites” was calculated independently for each replicate and experiment as the average log2 ratio of all corresponding evidence table entries. The replicate ratios were then averaged for the final ratio of the “phosphorylation site”.

### Protein-protein proximity assay (M-Track)

M-track protein protein proximity analysis (Brezovich et al., 2015; Zuzuarregui et al., 2012) was carried out using MES lysis buffer (50 mM MES/NaOH pH 6.5, 150 mM NaOH, 1% Triton, 1 mM EDTA, Complete EDTA free (Roche)). Cell extracts were prepared by glass bead lysis using a Fast Prep 24 instrument (MP Biomedicals) with the following settings: 1 x 45 s, power level 6.5. Immunoprecipitation of prey proteins was achieved using anti-HA-magnetic beads (Pierce). Washing steps were carried out using a MES-wash buffer (50 mM MES/NaOH pH 6.5, 150 mM NaOH, 1 mM EDTA). Prey proteins were eluted by heat incubation in 2 x urea sample buffer. Each sample was mixed with 4 μl of a BY4741 lysate (protein concentration 1 mg/ml in urea sample buffer) before loading in order to improve western blot transfer efficiency rates.

Eluates were analyzed by western blotting. Histone H3 Lysine 9 trimethylation (me3K9H3) of Rck2-protA-H3-HA was visualized using an antibody recognizing me3K9H3 (1:2000 dilution in 1% yeast extract (YE) in PBS-T, Novus #NBP1–30141). Membranes were incubated with primary antibody for 1◻h at 4◻°C, followed by 1◻h incubation at 4◻°C with HRP-conjugated goat anti-mouse (1:5000 dilution in 1% YE in PBS-T, BioRad #170–6516) secondary antibody. No washing steps were performed between primary and secondary antibody incubation. Loading was controlled using an antibody recognizing HA (1:5000 in PBS-T, 12CA5). PicoECL (Thermo Scientific) was used for enhanced chemiluminescent (ECL) detection.

### Phospho-shift assay

Cell lysis was carried out in MES-lysis buffer (50 mM MES/NaOH pH 6.5, 150 mM NaOH, 1% Triton, 1 mM EDTA, Complete EDTA free (Roche)) by beating using a Fast Prep 24 instrument (MP Biomedicals) with the following settings: 3 x 45 s, power level 6.5. Cleared protein extracts were resolved in SDS-PAGE loading buffer (62.5 mM Tris–HCl (pH 6.8), 8 M urea, 2% (w/v) SDS, 0.05% (w/v) bromophenol blue, 10% (v/v) glycerol, 5% (v/v) β-mercaptoethanol) and incubated at 95 °C for 1 min. For Mn^2+^-Phos-tag SDS-PAGE, 25 μM of Phos-tag-AAL (Wako) and 50 μM of MnCl_2_ were added to the separating gel before polymerization. The pH of the separation gel was adjusted to 8 for Mn^2+^-Phos-tag SDS-PAGE and 8.8 for standard SDS-PAGE. Gel mobility shifts were visualized by western blot using an antibody recognizing Flag (M2, Sigma-Aldrich). For visualization of blue protein bands from the prestained protein ladder the anti-BLUE antibody (Anti-BLUE, 2D2-F11, Millipore) was used (Schuchner et al., 2016). Each sample was mixed with 4 μl of a HeLa lysate (protein concentration 1 mg/ml in urea sample buffer) before loading in order to improve western blot transfer efficiency rates.

### Prediction of LxxIxE SLiM binding motif

Rts1 SLiM binding motifs were identified by first searching protein sequences with the regular expression pattern “([LFM]..[IVLCW].E)|([LFMYWIC]..[IVL].E)”. Matches had to be followed by an acidic patch, which was defined as at least two occurrences of the amino acids “D” and “E” in the subsequent six positions after the SLiM motif. To analyze whether putative SLiM motifs lie within a disordered region we utilized the SPOT-Disorder algorithm (Hanson et al., 2017).

## Acknowledgements

We would like to thank Dorothea Anrather for general MS-related support, and the VBCF for providing the MS instrument pool. This work was supported by the FWF Austrian Science Fund (grant numbers W1261 and F34). MH and WR were supported by the FWF Special Research Program F70.

## Authors’ contribution

DH and WR conceptualized the study. DH, EO, MH, GA and WR designed experiments. DH, GG, JV and WR performed experiments. DH and NR analyzed the data. DH, NR, and WR wrote the paper. All authors edited the text. All authors read and approved the final manuscript.

